# A minimalist approach to protein identification

**DOI:** 10.1101/187013

**Authors:** G. Sampath

## Abstract

Computations on proteome sequence databases show that most proteins can be identified from a protein’s isoelectric point (IEP) and digitized linear sequence volume (equal to the total volume of its residues). This is illustrated with four proteomes: *H. pylori* (1553 proteins), *E. coli* (4306 proteins), *S. cerevisiae* (6721 proteins), and *H. sapiens* (20207 proteins); the identification rate exceeds 90% in all four cases for appropriate parameter values. IEP can be obtained with 1-d gel electrophoresis (GE), whose accuracy is better than 0.01. Linear protein sequence volumes of unbroken proteins can be obtained with a sub-nanometer diameter nanopore that can measure residue volume with a resolution of 0.07-0.1 nm^3^ (Kennedy et al., *Nature Nanotech*., 2016, **11**, 968-976; Dong et al., *ACS Nano*, 2017, doi: 10.1021/acsnano.6b08452); the blockade current due to a translocating protein is roughly proportional to the volume it excludes in the pore. There is no need to identify any of the residues. More than 90% of all the proteins have estimated translocation times higher than 1 μs, which is within the time resolution of available detectors. This is a minimalist proteolysis-free GE-and nanopore-based single-molecule approach requires very small samples, is non-destructive (the sample can be recovered for reuse), and can be translated with currently available technology into a portable device for possible use in the field, an academic lab, or a pre-screening step preceding conventional mass spectrometry.

## 1. Introduction

In multidimensional identification of proteins [1-3], predominantly the domain of mass spectrometry (MS), multiple physical/chemical properties (electric charge/mobility, hydrophobicity, isoelectric point (IEP), mass (molecular weight), diffusion constant, etc.) of the full protein or peptides obtained from it by proteolysis are measured, and the results matched with corresponding values for known proteins. Recently a nanopore has been used to obtain a five-dimensional identifier for a known protein [4], and also as a spectrometer to measure peptide mass spectra [5]. The complexity of the identification space varies with the approach. Thus isoelectric focusing locates proteins (or peptides) in 1-dimensional IEP space [6], 2-d gel electrophoresis (GE) locates them in IEP × mass [7], LC-MS (liquid chromatography followed by MS) locates ionized peptides in hydrophobicity × peptide mass-to-charge ratio [8]. Nanopore-based 5-d identification locates a protein in the space defined by shape × (folded) volume × charge × rotational diffusion coefficient × dipole moment [4]. Experiment is usually followed by search through a proteome sequence database [9].

The present work considers a characteristic of proteins that has not been previously used for protein identification, namely the linear volume of a protein, which is equal to the sum of the spatial volumes of the residues of the protein stretched end to end on a line. In combination with IEP this defines the identification space IEP × linear-sequence-volume. Such an approach can identify a vast majority of proteins in a proteome; this is illustrated computationally with the proteomes of *H. pylori*, *E. coli*, *S. cerevisiae*, and *H. sapiens*, with the identification rate exceeding 90%. This computational model can be translated into practice by first obtaining IEP via 1-d gel electrophoresis, which can separate proteins with an accuracy better than 0.01, then measuring the linear sequence volumes of separated proteins with a nanopore as they translocate single file linearly stretched out through the pore [10-12]. No residues are identified in the process. Volume resolution in the range 0.07-1.0 nm^3^ is possible with a sub-nanometer diameter nanopore [13,14]. o proteolysis is involved so the sample can be recovered and reused any number of times. The method proposed here can be translated with available technology into a hand-held device that can potentially be used in field studies, in an academic laboratory, or in a pre-screening step before conventional MS.

## 2. Methods

Complete sets of protein sequences were downloaded from the Uniprot website http://www.uniprot.org for the following four proteomes: the gut bacterium *Helicobacter pylori* (Uniprot id UP000000210, 1553 sequences), the pathogen *Escherichia coli* (Uniprot id UP000000562; 4306 sequences), baker’s yeast *Saccaromyces cerevisiae* (Uniprot id UP000004932; 6721 sequences), and human *Homo sapiens* (Uniprot id UP000005640; 20207 curated sequences). For the four proteomes pI (see below) values and digitized values of linear sequence volumes were calculated for different parameter values.

Let the set of amino acids be **AA** = [A, R, N, D, C, Q, E, G, H, I, L, K, M, F, P, S, T, W, Y, V] where AA_i_ is the i-th amino acid, 1 ≤ i ≤ 20. The mass array is **AAmass** = [71.04, 156.1, 114.04, 115.03, 103.01, 129.04, 128.06, 57.02, 137.06, 113.08, 113.08, 128.09, 131.04, 147.07, 97.05, 87.03, 101.05, 186.08, 163.06, 99.07] and is based on [15]; all values are in daltons. The mean volume array **AAvol** = [87.8, 188.2, 120.1, 115.4, 105.4, 145.1, 140.9, 59.9, 156.3, 166.1, 168, 172.7, 165.2, 189.7, 123.3, 91.7, 118.3, 227.9, 191.2, 138.8] and is based on [16]; all values are in 10^-3^ nm^3^.

### Calculating the charge carried by a protein

Let **P** = p_0_ p_1_ … p_N-1_ be the primary sequence of a protein of length N from a proteome with M proteins, where the p_i_’s are residues from **AA**. The charge carried by **P** at a given pH, C(**P**, pH), is calculated with the Henderson-Hasselbalch equation [17].

### Calculating the isoelectric point (pI value) of a protein

The pI value for a protein **P** is the pH value at which C(**P**, pH) = 0. The following procedure, which assumes that the pI value is a unimodal function of pH, calculates pI for each protein in a proteome:

**Procedure** Calculate-pI-values-for-all-proteins-in-proteome

**for** each protein **P**_i_, 0 ≤ i < M:

Set pH = pH_last = 0.

**while** (pH < 14 and not found pI):

Calculate the charge C(**P**_i_, pH) carried by **P**_i_.

**if** C(**P**_i_, pH) C(**P**_i_, pH_last) < 0 set pI_i_ to pH_last and exit loop

**else** set pH_last to pH; set pH to pH+increment

**end if**

**end while**

return pH_last as the pI value for the protein

**end for**

### Calculating the digitized linear sequence volume of a protein

Let I(p_i_) be the position of p_i_ in **AA**. The (mean) volume of a linearly stretched protein sequence of length L is

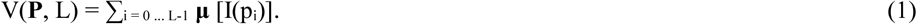

which is the sum of the individual residue volumes. The mean volumes of the 20 standard amino acids are given by the array **AAvol** above. The digitized value of V(**P**, L) based on a volume resolution of D is

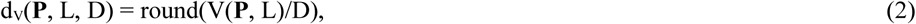

where the argument is rounded up to the next integer if the fractional part ≥ 0.5 and truncated otherwise.

### Protein identification in a proteome

Based on the above development, protein i is uniquely characterized by the pair (IEP_i_, d_V_(**P**_i_, L, D)) if its IEP and/or d_V_ value differs from the IEP and d_V_ values of every other protein j in the proteome by given amounts IEPdiff and dVdiff. The following procedure was used to identify protein **P**_i_ from its (**IEP_i_**, d_V_(**P**_i_, L, D)) pair as calculated above:

**Procedure** Identify_Protein(IEPdiff, dVdiff)

1. For each protein **P**_i_ obtain its IEP and digitized linear sequence volume as (IEP_i_, d_V_(**P**_i_, L, D)).
2. For every protein j ≠ i compute |IEP_i_, - IEP_i_| and |d_V_(**P**_i_, L, D) – d_V_(**P**_j_, L, D)|. If for all j ≠ i |IEP_i_ - IEP_i_| ≥ IEPdiff or |d_V_(**P**_i_, L, D) – d_V_(**P**_j_, L, D)| ≥ dVdiff mark this protein i as unique.
3. Output the sequence number and protein id in the proteome for each unique protein and the total number of unique proteins..

Notice that this is just a matter of identifying a protein; no sequencing is done so no residues are identified here.

## 3. Results

Table 1 (displayed at the end after References) shows the percentage of proteins that are identified uniquely vs the difference in the digital volume dVdiff between two protein sequences for volume resolution D ϵ {50, 70, 100} and IEPdiff ϵ {0.05, 0.1} for each of the four proteomes.

**Table 1.** Percentage of proteins identified in four different proteomes vs difference in digital volumes (dVdiff) between two protein sequences for three values of linear sequence volume resolution D ϵ {50, 70, 100} 10^−3^ nm^3^ of the digital resolution used in measuring sequence volume via the blockade current, and two values of pi (IEP) difference in gel electrophoresis: IEPdiff ϵ {0.05, 0.1}. (Percentages for a projected (that is, not yet realized) digital volume resolution of D = 50.0 are shown in italics.)

|  | Percentage of proteins identified <sup>(a)</sup> |  |  |  |  |  |  |  |  |  |  |  |
| --- | --- | --- | --- | --- | --- | --- | --- | --- | --- | --- | --- | --- |
|  | *D = 50.0* |  | D = 70.0 |  | D = 100.0 |  | *D = 50.0* |  | D = 70.0 |  | D = 100.0 |  |
|  | *IEPdiff* |  | IEPdiff |  | IEPdiff |  | *IEPdiff* |  | IEPdiff |  | IEPdiff |  |
|  | 0.05 | 0.10 | 0.05 | 0.10 | 0.05 | 0.10 | 0.05 | 0.10 | 0.05 | 0.10 | 0.05 | 0.10 |
| **dVdiff** | *H. pylori* (1553 proteins) |  |  |  |  |  | *E. coli* (4306 proteins) |  |  |  |  |  |
| 1 | 99.23 | 97.68 | 98.71 | 97.17 | 97.81 | 95.36 | 96.21 | 93.89 | 95.29 | 91.76 | 93.89 | 89.55 |
| 2 | 97.42 | 94.27 | 96.01 | 91.69 | 95.36 | 88.99 | 91.62 | 85.28 | 88.27 | 79.96 | 84.86 | 73.8 |
| 3 | 96.01 | 90.73 | 94.91 | 87.31 | 90.99 | 81.65 | 87.04 | 77.59 | 82.68 | 70.97 | 76.08 | 61.68 |
| 4 | 94.46 | 87.25 | 92.21 | 83.26 | 89.18 | 77.59 | 82.51 | 70.74 | 76.36 | 62.17 | 69.74 | 53.18 |
| 5 | 92.79 | 84.22 | 89.89 | 79.27 | 87.06 | 73.92 | 78.68 | 64.82 | 71.74 | 56.18 | 64.14 | 46.31 |
| **dVdiff** | *S. cerevisiae* (6721 proteins) |  |  |  |  |  | *H. sapiens* (20207 proteins) |  |  |  |  |  |
| 1 | 94.91 | 92.95 | 93.86 | 91.01 | 92.52 | 88.77 | 92.84 | 87.24 | 90.41 | 82.92 | 86.79 | 77.32 |
| 2 | 89.7 | 84.24 | 87.13 | 79.82 | 83.62 | 74.16 | 81.27 | 68.73 | 75.4 | 60.52 | 67.92 | 50.63 |
| 3 | 85.51 | 77.32 | 81.71 | 71.22 | 76.61 | 62.97 | 72.05 | 55.95 | 64.35 | 46.24 | 55.23 | 36.54 |
| 4 | 81.64 | 70.82 | 76.63 | 63.34 | 70.18 | 54.19 | 64.39 | 46.3 | 55.71 | 36.93 | 45.64 | 27.72 |
| 5 | 78.23 | 65.42 | 72.16 | 57.0 | 64.66 | 47.21 | 58.01 | 39.28 | 48.56 | 30.21 | 38.71 | 22.24 |

## 4. Discussion

The following are some practical and computational considerations.

1. The computational model of protein identification described above can be implemented with the electrical current equivalent of residue volume. Recently a nanopore with diameter < 1 nm was shown to be capable of measuring residue volume with a resolution of 0.07 nm^3^ [13]. A similar pore, with a volume resolution of 0.1 nm^3^, has been used in identifying single amino acid substitutions in a protein [14]. Assuming for simplicity that linear sequence volume maps linearly to blockade current measured, a translocating protein is represented by the measured pair (*pI*, *digitized value of blockade current integral*).
2. In the computation of IEP above (Section 2), which is based on Lehninger’s method [17], corrections can be made for the influence that neighboring residues might have on the kA value of a residue; see [18] for a review of several schemes.
3. With a gel strip 28 cm long, an IEP accuracy of 0.05 can be obtained with slices 1 mm thick. With slices 0.4 mm thick the identification rate rises to the 95-98% range over all four proteomes.
4. The detector used to measure the blockade current in a nanopore must have adequate bandwidth. Currently the smallest time resolution available is ~1 μs (equivalent to a bandwidth of ~500 Khz) [19]. Recent work on synthetic nanopores 6-8 nm thick show that proteins with molecular weights under 30 kD have average translocation times around ~2.5 μs [20]. Using this as a baseline, the results of Section 3 show that over 90% of the proteins in the four proteomes considered here can be detected during their passage through a pore 6-8 nm thick and their sequence volumes measured.
5. Proteins tend to be weakly charged so they depend primarily on diffusion to enter the pore. One way to enable entry is to attach a charged carrier molecule like DNA [21]. Another is to treat proteins with thiol and sodium dodecyl sulphate (SDS). This unfolds and straightens out the protein and gives it a uniform negative charge along its length [22]; a potential of ~100 mV across the pore membrane draws the protein into the pore. As described in [14] the SDS gets stripped out at the entrance to the narrow pore, leaving the denatured straightened molecule to translocate through the pore by a combination of diffusion and electrophoresis. Passage of the protein is detected as a current blockade pulse whose integral over the duration of translocation is roughly proportional to the linear sequence volume of the protein. An A/D converter outputs the digitized value of this integral, which is then used, along with the pI value obtained from 1-d gel electrophoresis, for identification.
6. When a third dimension of molecular weight is added, computations in IEP × mass × Linear-sequence-volume space show that the identification rate increase is marginal. This suggests that 1-D GE is sufficient and 2-D GE is not necessary, so SDS-PAGE (and the associated complexity of spot picking [23]) can be bypassed. Output from the 1-D GE step (proteins from gel slices eluted and treated with SDS) can be input directly to a nanopore.
7. The method proposed here keeps proteins intact, they can be recovered from the *trans* compartment of the electrolytic cell containing the nanopore. In contrast MS requires proteins to be broken into peptides, which then have to be ionized (using a method like electrospray ionization (ESI)) before they enter the spectrometer.
8. This is a minimalist approach to protein identification. Its objective is the development of a low-cost alternative to high-end methods like LC-MS (the technology underlying precision proteomics [24]) in the form of a hand-held device that can be used in the field or in an academic lab, or in a pre-screening step preceding MS.

## Supplementary Information

Four files containing protein id, length, digital volume, total sequence volume, molecular weight, pI (IEP) value, and SDS fraction for each protein, one for each of *H. pylori* (1553 proteins), *E. coli* (4306 proteins), *S. cerevisiae* (6721 proteins), and *H. sapiens* (Uniprot id UP000005640; 20207 proteins).

